# BATCAVE: Calling somatic mutations with a tumor- and site-specific prior

**DOI:** 10.1101/798348

**Authors:** Brian K. Mannakee, Ryan N. Gutenkunst

## Abstract

Detecting somatic mutations withins tumors is key to understanding treatment resistance, patient prognosis, and tumor evolution. Mutations at low allelic frequency, those present in only a small portion of tumor cells, are particularly difficult to detect. Many algorithms have been developed to detect such mutations, but none models a key aspect of tumor biology. Namely, every tumor has its own profile of mutation types that it tends to generate. We present BATCAVE (Bayesian Analysis Tools for Context-Aware Variant Evaluation), an algorithm that first learns the individual tumor mutational profile and mutation rate then uses them in a prior for evaluating potential mutations. We also present an R implementation of the algorithm, built on the popular caller MuTect. Using simulations, we show that adding the BATCAVE algorithm to MuTect improves variant detection. It also improves the calibration of posterior probabilities, enabling more principled tradeoff between precision and recall. We also show that BATCAVE performs well on real data. Our implementation is computationally inexpensive and straightforward to incorporate into existing MuTect pipelines. More broadly, the algorithm can be added to other variant callers, and it can be extended to include additional biological features that affect mutation generation.

## Introduction

Cancer develops through the accumulation of somatic mutations and clonal selection of cells with mutations that confer an advantage. Understanding the evolutionary history of a tumor, including the mutations that drive its growth, the genetic diversity within it, and the accumulation of new mutations, requires accurate variant identification, particularly at low variant allele frequency [1, 2, 3, 4]. Accurate variant calling is also critical for optimizing the treatment of individual patients’ disease [5, 6, 7, 8, 9]. Low frequency mutations challenge current variant calling methods, because their signature in the data is difficult to distinguish from the noise introduced by Next Generation Sequencing (NGS), and this challenge increases with sequencing depth.

Many methods have been developed for calling somatic mutations from NGS data. The earliest widely used somatic variant callers developed specifically for tumors, MuTect1 [10] and Varscan2 [11], used a combination of heuristic filtering and a model of sequencing errors to identify and score potential variants and set a threshold score designed to balance sensitivity and specificity. Subsequent research gave rise to a number of alternate strategies, including haplotype-based calling [12], joint genotype analysis (SomaticSniper [13], JointSNVMix2 [14], Seurat [15], CaVEMan [16], and MuClone [17]), allele frequency-based analysis (Strelka [18], LoFreq [19], EBCall [20], deepSNV [21], LoLoPicker [22], and MuSE [23]), and ensemble and deep learning methods (MutationSeq [5], BAYSIC [24], SomaticSeq [25], and SNooPer [26]). These methods vary in their complexity and specific focus. But they all implicitly or explicitly assume that the rate of mutation is uniform across the genome.

The mutational processes that generate single nucleotide variants in tumors do not act uniformly across the genome. If fact, even the processes of spontaneous mutation that are active in all somatic tissues depend sensitively on local nucleotide context [27, 28, 29]. Additional mutational processes are active in tumors, due to mutagen exposure or defects in DNA maintenance and repair, and these processes are also sensitive to local nucleotide context [30, 31, 32, 33, 34]. The specific mutational processes active in a particular tumor generate its unique mutation profile, and differences within and between tumor types are pronounced [35, 36, 37, 38, 39]. For example, the mutation profiles differ substantially among the three breast tumors illustrated in Figure 1B-D.

**Figure 1:**
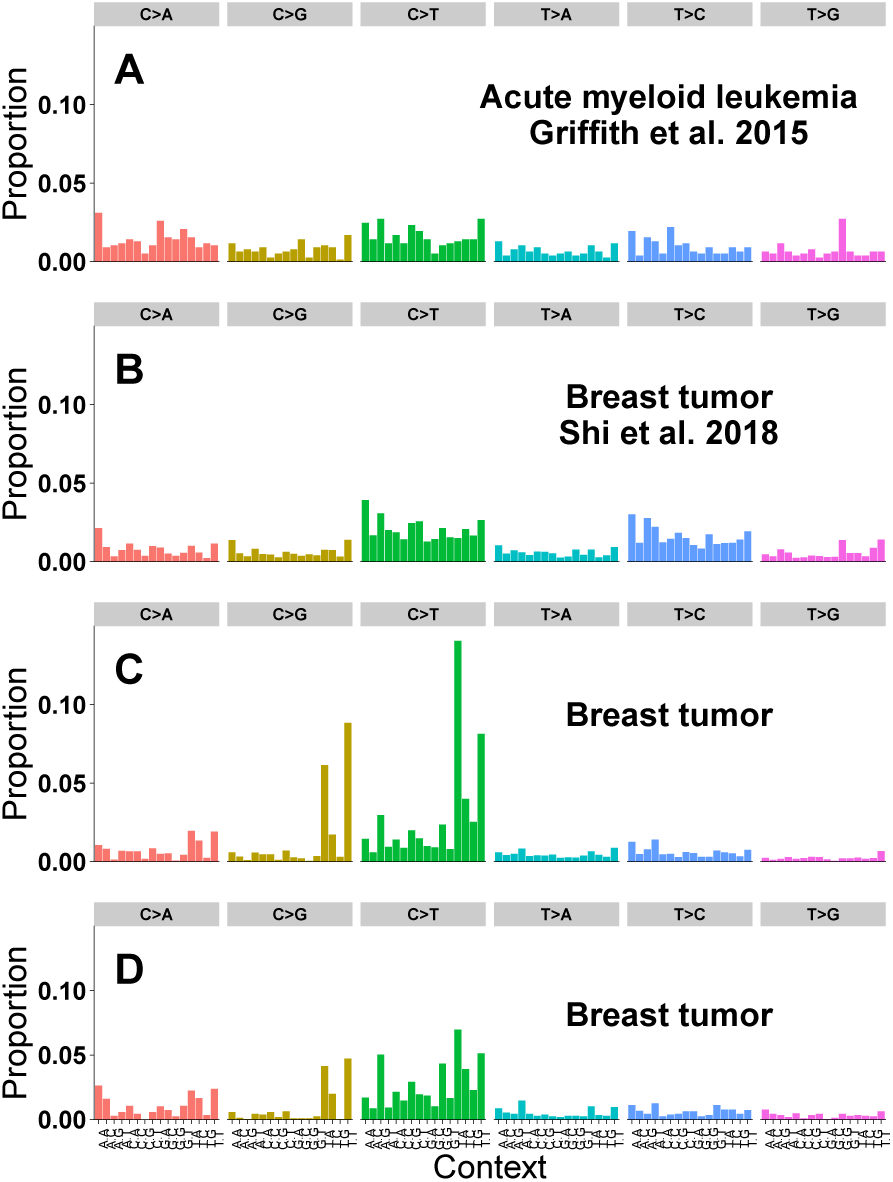
Real tumor mutation profiles. In each panel, the x-axis corresponds to each of the 96 possible mutation types, and the y-axis is the proportion of total mutations of each type. (A) The observed mutation profile of an acute myeloid leukemia used in our real data analysis [40]. (B) The observed mutation profile of a breast tumor used in our real data analysis [4]. (C)&(D) The observed mutation profiles of two additional breast tumors [41].

Here we present an enhanced variant-calling algorithm that uses the biology of each individual tumor’s mutation profile to improve identification of low allelic frequency mutations. Our BATCAVE algorithm first estimates the tumor’s mutation profile and mutation rate using high-confidence variants and then uses them as a prior when calling other variants. Our R implementation of the algorithm, batcaver, takes output from the MuTect variant caller as input and returns the posterior probability that a site is variant for every site observed by MuTect. Using both simulated and real data, we show that the addition of a mutation profile prior to MuTect produces a superior variant caller. Our algorithm is simple and computationally inexpensive, and it can be integrated into numerous other variant callers. Broad adoption of our approach will enable more confident study of low allelic frequency mutations in tumors in both research and clinical settings.

## MATERIALS AND METHODS

### Somatic variant calling probability model

At every site in the genome with non-zero coverage, Next Generation Sequencing produces a vector **x** = ({*b*_*i*_}, {*q*_*i*_}), *i* = 1 …d of base calls *b* and their associated quality scores *q*, where d is local read depth. Variant callers use the data **x** to choose between competing hypotheses:

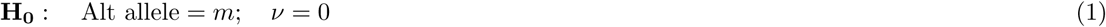

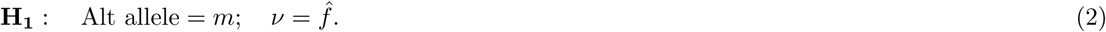

Here *m* is any of the 3 possible alternate non-reference bases and *ν* is the variant allele frequency. The maximum likelihood estimate of *ν* is simply 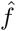, the number of variant reads divided by the local read depth. The posterior probability of a given hypothesis, P(*m, ν*), is the product of the likelihood of the data given that hypothesis and the prior probability of that hypothesis. Assuming that reads are independent, this is

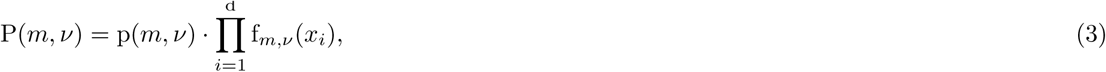

where f_*m,ν*_ (*x*_*i*_) is the probability model for reads, and *p*(*m, ν*) is the prior.

Assuming that the identity of the alternate allele and its allele frequency are independent and that *ν* is uniformly distributed, Eq. 3 becomes

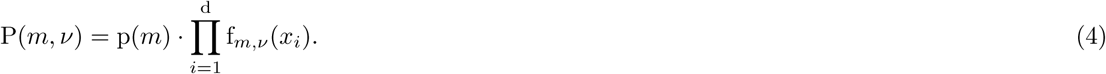

The focus of BATCAVE is to provide a tumor- and site-specific estimate of the prior probability of mutation p(*m*).

### Site-specific prior probability of mutation

The probability that we have denoted p(*m*) in Eq. 4 is more precisely the joint probability that a mutation has occurred *M* and that it was to allele *m*, which we denote p(*m, M*). But p(*m, M*) is not uniform across the genome. Rather it depends on the local genomic context *C*, so its full form is p(*m, M*|*C*) [42]. Assuming that *m* and *M* are independent conditional on the genomic context, p(*m, M | C*) = p(*m | C*)p(*M | C*), which we can use Bayes’ theorem to further decompose as

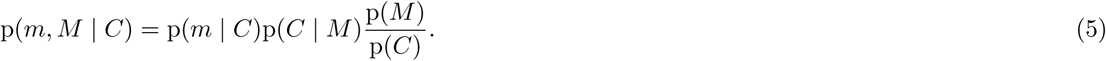

We next show how to estimate the quantities in Eq. 5.

### Estimation of the mutation profile

Many aspects of genomic architecture can affect the somatic mutation rate at multiple scales [42]. Here we focus on a small-scale feature, the trinucleotide context, which is known to strongly affect the prior probability of single-nucleotide mutation [27, 28, 29]. The trinucleotide context of a genomic site consists of the identity of the reference base and the 3’ and 5’ flanking bases. Folding the central base to the pyrimidines, there are two possible bases at the focal site, and there are four possible bases 3’ and 5’ of the focal site, yielding 2 *⋅* 4 *⋅* 4 possible tri-nucleotide contexts *C*. At the focal site, a mutation *m* can be to any of three alternate alleles. Indexing by the *c* = {1 … 32} contexts and by the *m* = {1 … 3} alternate bases, we have 96 possible substitution types *S*_*m,c*_. Eq. 5 is then

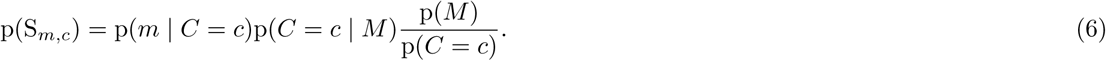

The first two terms on the right-hand side can be estimated from the observed mutation profile (Fig. 1).

We model the observed mutation profile S as multinomial with parameter ***π*** = *π*_*m,c*_. Each element of ***π*** represents the expected proportion of mutations that are to allele *m* and in context *c*. In a tumor with many high-confidence observed mutations, ***π*** could be estimated directly from the observed mutation profile *S*. But in practice many entries in ***π*** would then have zero weight. We thus model the distribution of S as Dirichlet-multinomial with pseudo-count hyper-parameter ***α***,

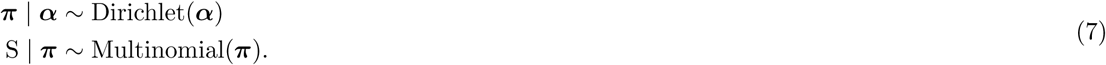

In BATCAVE we use the symmetric non-informative hyper-parameter ***α*** = **1**, so a priori mutation is equally likely to any allele and in any context.

To estimate ***π***, we identify a subset of high confidence variants, based on an initial calculation of their likelihood given the data. These are variants for which the evidence in the read data overwhelms any reasonable value of the site-specific prior probability of mutation. Let D be the set of high confidence variant calls, which we define as those having posterior odds greater than 10 to 1 without the site-specific prior, and s *∈* D be the substitution type of each mutation in D. The posterior distribution of ***π*** is then p(***π*** | D) *∼* Dirichlet(***α′***) where

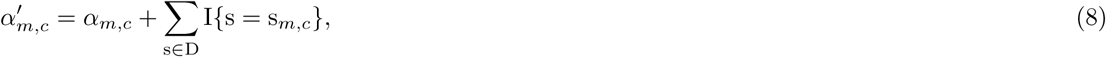

and *I* is the indicator function. Returning to Eq. 6, given that a mutation has occurred, the posterior probability it occurred in context *c* is

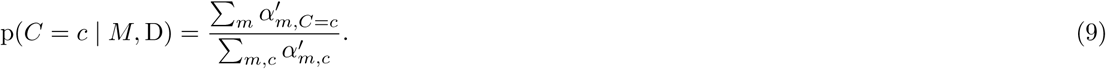

The posterior probability of mutation to allele *m* given that a mutation has occurred in context *C* = *c* is then

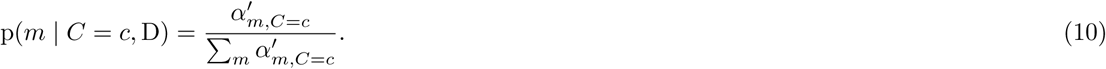

The prior probability of each particular trinucleotide context *p*(*C* = *c*) is computed simply as the proportion of sequenced trinucleotide contexts that have context *c*. The R implementation of BATCAVE ships with pre-computed tables for both human whole exomes and whole genomes.

### Estimation of the mutation rate

The final piece of Eq. 6 is *p*(*M*), the prior probability of mutation, which we specify as the per-base per-division mutation rate *µ*. In an exponentially growing and neutrally evolving tumor, branching process calculations [3] show that the expected total number of mutations M_tot_ between two allele frequencies (*f*_*min*_,*f*_*max*_) is

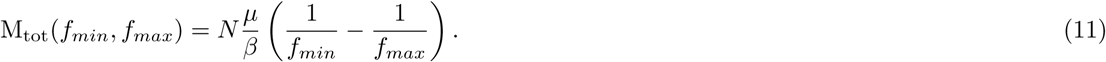

The number of bases *N* is 3 ⋅ 10^9^ for a whole genome and 3 ⋅ 10^7^ for a whole exome. The quantity *µ/β* is the effective mutation rate, where *β* is the fraction of cell divisions that lead to two surviving lineages. We make the simplifying assumption that there is no cell death (*β* = 1), so we somewhat over-estimate *µ*. We then estimate *µ* by counting observed high-confidence mutations between allele frequencies *f*_*min*_ and *f*_*max*_. We set *f*_*max*_ to be the largest allele frequency in D, but we must choose *f*_*min*_ conservatively, depending on sequencing depth. In the R implementation of BATCAVE, *f*_*min*_ is a free parameter. For this paper, we set *f*_*min*_ = 0.05, because we are working at high depth.

### Likelihood function

The current implementation of BATCAVE builds on MuTect, because MuTect reports the log ratio of the likelihood functions for the null and alternative hypotheses (Eq. 1) as TLOD (MuTect1) or t lod fstar (MuTect2). We used MuTect 1.1.7 for all analyses in this paper, so we have

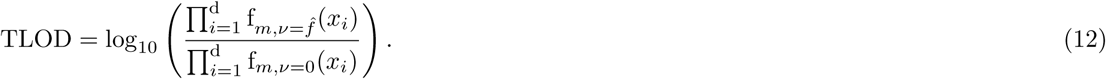

The log posterior odds is the log likelihood ratio (TLOD) plus the log prior odds, so the posterior odds in favor of the alternate hypothesis for a given substitution type is

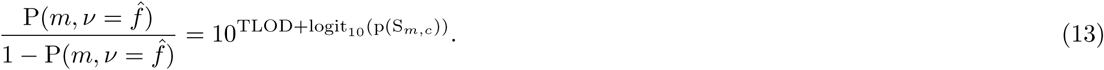

Here p(S_*m,c*_) is the prior probability of a substitution of type S_*m,c*_, as described in Eq. 6 and specified in Eq. 9-11. When comparing our posterior odds to those of MuTect, we assume a uniform per-base probability of mutation of 3 *⋅* 10^*−*6^ [10], so

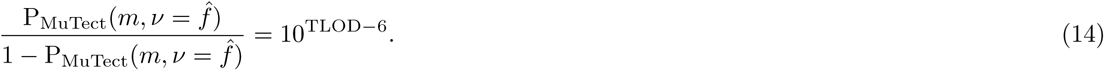

### Implementation

We have implemented the BATCAVE algorithm as an R package `batcaver`. The package leverages the Bioconductor packages BSgenome [43], GenomicAlignments [44], VariantAnnotation [45], and SomaticSignatures [46] for fast and memory-efficient variant annotation and genomic context identification. Reference sequences are specified as BSgenome objects, allowing efficient access to genomic context information.

### Tumor simulations

We used a neutral branching process with no death and *µ* = 3 ⋅ 10^*−*6^ to simulate realistic distributions of mutation frequencies. Tumors were simulated with three different mutation profiles composed of COSMIC mutation signatures (version 2) [47]. Each simulated profile includes COSMIC signature 1, which is found in nearly all tumors and is associated with spontaneous cytosine deamination. The “Concentrated” profile (Fig. 2A) is an equal combination of COSMIC signatures 1, 7, and 11, which has a large percentage of C *>* T substitutions such as are often seen in cancers caused by UV exposure [48]. The “Intermediate” profile (Fig. 2B) is an equal combination of COSMIC signatures 1, 4, and 5, which has been associated with tobacco carcinogens and is representative of some lung cancers [48]. The “Diffuse” profile (Fig. 2C) is an equal combination of COSMIC signatures 1, 3, and 5, which has been associated with inactivating germline mutations in the BRCA1/2 genes leading to a deficiency in DNA double strand break repair [32]. Simulated variants were sampled from a combination of the Cancer Genome Atlas (TCGA) and Pan-Cancer Analysis of Whole Genomes (PCAWG) databases, which include mutations found in all types of cancer. Whole genome (100X depth) and whole exome (500X depth) reads were simulated from the GRCh38 reference genome using VarSim [49] and aligned with BWA [50], both with default parameters. Variants were inserted to create tumors with BAMSurgeon with default parameters [51] and called with MuTect 1.1.7 [10] with the following parameters:

**Figure 2:**
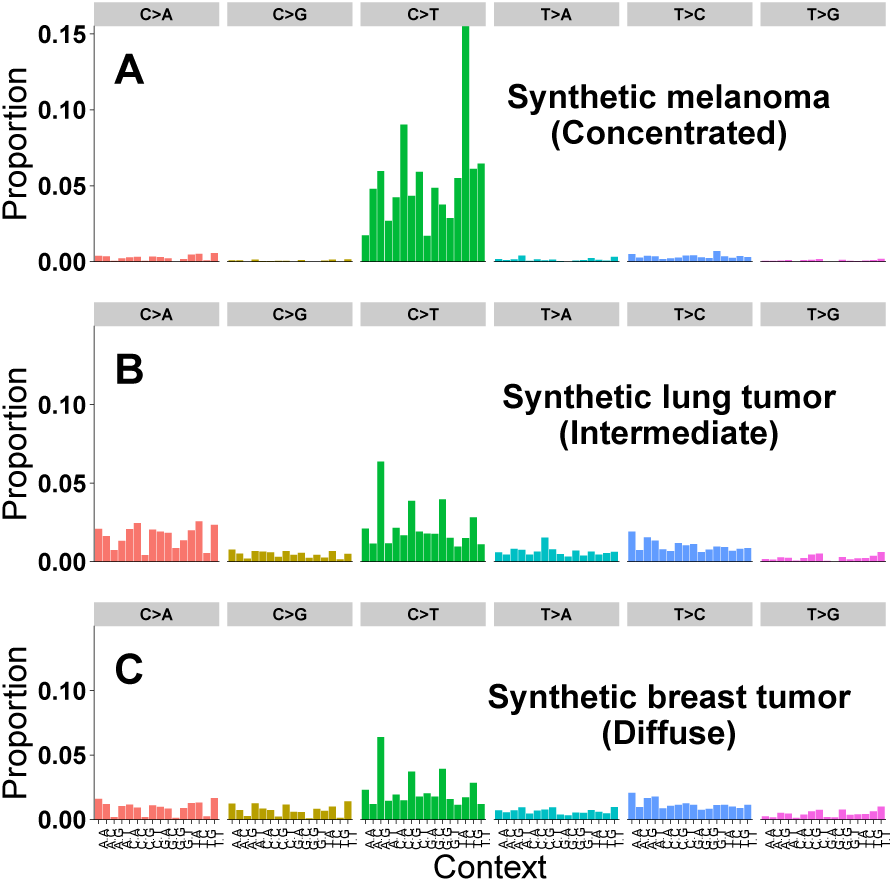
Simulated tumor mutation profiles. As in Fig. 1, in each panel the x-axis corresponds to each of the 96 possible mutation types, and the y-axis is the proportion of total mutations of each type. (A) A mutation profile used for simulating tumors, made up of equal proportions of COSMIC mutation signatures 1, 7, & 11. (B) Equal proportions of signatures 1, 4, & 5. (C) Equal proportions of signatures 1, 3, & 5.

~~~
java -Xmx24g -jar $MUTECT JAR --analysis type MuTect --reference sequence $ref path --dbsnp $db snp
--enable extended output --fraction contamination 0.00 --tumor f pretest 0.00 --initial tumor lod −10.00
--required maximum alt allele mapping quality score 1 --input file:normal $tmp normal --input file:tumor
$tmp tumor --out $out path/$chr.txt --coverage file $out path/$chr.cov.
~~~

Variants identified by MuTect are labelled as to whether they pass all filters, fail to pass only the the evidence threshold tlod f star filter, or fail to pass any other filter. Variants that passed all filters or failed only tlod f star were then passed to BATCAVE for prior estimation and rescoring.

### Calibration metric

To quantify the difference in calibration between MuTect and BATCAVE, we used the Integrated Calibration Index [52]. Briefly, a loess-smoothed regression was fit by regressing the binary (True=1, False=0) true variant classification against the reported posterior probability for both MuTect and BATCAVE. For a perfectly calibrated caller, the regression fit would be the diagonal line *y* = *x*. The Integrated Calibration Index is a weighted average of the absolute distance between the calibration curve and the diagonal line of perfect calibration.

### Real data

We analyzed two real data sets, one from an acute myeloid leukemia (AML) [40] and one from a multi-region sequencing experiment in breast cancer [4]. We downloaded the normal and primary whole-genome AML tumor bam files from dbGaP accession number phs000159.v8.p4. Griffith et al. generated a platinum set of variant calls for this tumor [40], which we used for our true positive dataset. We downloaded the normal and tumor whole-exome breast cancer bam files from NCBI Sequence Read Archive accession SRP070662. Shi et al. generated a gold set of variant calls for each tumor region sequenced [4], which we used for our true positive dataset. For these multi-region data, we ran BATCAVE separately on each sequenced region and combined results to generate precision-recall curves. We called variants using Mutect 1.1.7 as in our simulations, except that both these data sets were originally aligned to GRCr37, so we used that reference.

## RESULTS

We implemented BATCAVE as a post-call variant evaluation algorithm to be used with MuTect (Versions 1.1.7 or *>*2.0) [10]. BATCAVE extracts the log-likelihood ratio for each potential variant site from the MuTect output, and then it uses that ratio to separate the potential sites into high and low confidence groups. The mutation profile and mutation rate are estimated from the high confidence sites, and the posterior probability of mutation is then recomputed for all sites. The BATCAVE algorithm is inexpensive, processing 22,000 variants per second on a typical desktop computer, which corresponds to roughly 100 seconds to process a 500X exome and 2,000 seconds for a 100X whole genome.

To test the performance of BATCAVE, we generated six different tumor/normal pairs, corresponding to 100X whole genomes and 500X whole exomes for three different mutation profiles. The three mutation profiles were chosen to resemble a melanoma (concentrated), a lung cancer (intermediate), and a BRCA-driven breast cancer (diffuse) (Fig. 2). We also tested BATCAVE using two real cancer data sets, a whole-genome Acute Myeloid Leukemia (AML) [40] and a whole-exome multi-region breast cancer [4]. In both, deep sequencing and variant validation were performed with the specific purpose of evaluating tumor variant calling pipelines. Because our focus is on evaluating the statistical calling model, we computed all test metrics using only those potential variants that passed MuTect’s heuristic filters and entered the statistical model.

### Tests using simulated data

To improve variant identification, the context-dependent prior probability of mutation must converge to an accurate representation of the data generating distribution within the set of high-confidence mutations. When applied to simulated data, the prior converged within a few hundred mutations (Fig. 3). For comparison, in our simulated data sets the number of high-confidence mutations ranged between 1,500 and 5,000, and in the real AML we test on it is over 17,000 [40].

**Figure 3:**
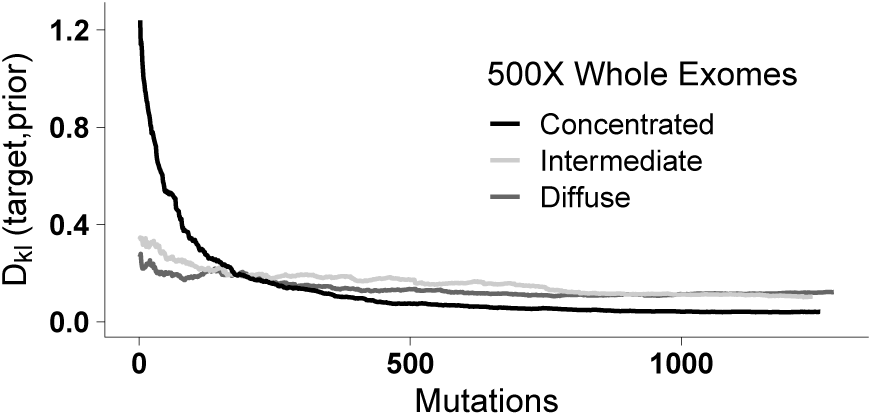
Convergence of the mutational prior to the data generating distribution. Plotted is the Kullback-Leibler divergence between the simulated and estimated profiles versus number of incorporated mutations for whole exomes. Convergence for whole genomes is similar.

We assessed classification performance using the areas under both the receiver operating characteristic and the precision-recall curves, because the classes are unbalanced (approximately 5 to 1 ratio of false to true variants in our simulated data). By both metrics BATCAVE outperforms MuTect (Fig. 4A&B, Fig. S1A&B, and Table 1). The extent of the performance difference is dependent on both the sequencing depth and the concentration of the mutation profile. Deeper sequencing and more concentrated mutation profiles increase the performance advantage of BATCAVE.

**Table 1:** Variant calling metrics for all data sets.

| Scenario | Mutation profile | $\mu$<br>(estimated) | AUROC | | AUPRC | | ICI | |
| --- | --- | --- | --- | --- | --- | --- | --- | --- |
|  |  |  | MuTect | BATCAVE | MuTect | BATCAVE | MuTect | BATCAVE |
| 100X whole genome | Concentrated | 3.6e-7 | .987 | .993 | .972 | .975 | .117 | .287 |
| 100X whole genome | Intermediate | 3.2e-7 | .987 | .989 | .972 | .973 | .118 | .214 |
| 100X whole genome | Diffuse | 3.2e-7 | .988 | .989 | .971 | .973 | .120 | .219 |
| 500X whole exome | Concentrated | 3.6e-7 | .848 | .929 | .674 | .758 | .138 | .109 |
| 500X whole exome | Intermediate | 3.6e-7 | .847 | .881 | .677 | .706 | .108 | .112 |
| 500X whole exome | Diffuse | 3.6e-7 | .850 | .873 | .676 | .698 | .105 | .116 |
| real AML [40] | Actual | 3.6e-8 | — | — | .995 | .996 | — | — |
| real breast [4] | Actual | 3.6e-8 | — | — | .972 | .972 | — | — |

$\mu$ = per-base mutation rate, AUROC/AUPRC = Area Under Receiver Operating Characteristic / Precision-Recall Curve, ICI = Integrated Calibration Index. Smaller values of ICI are superior.

**Figure 4:**
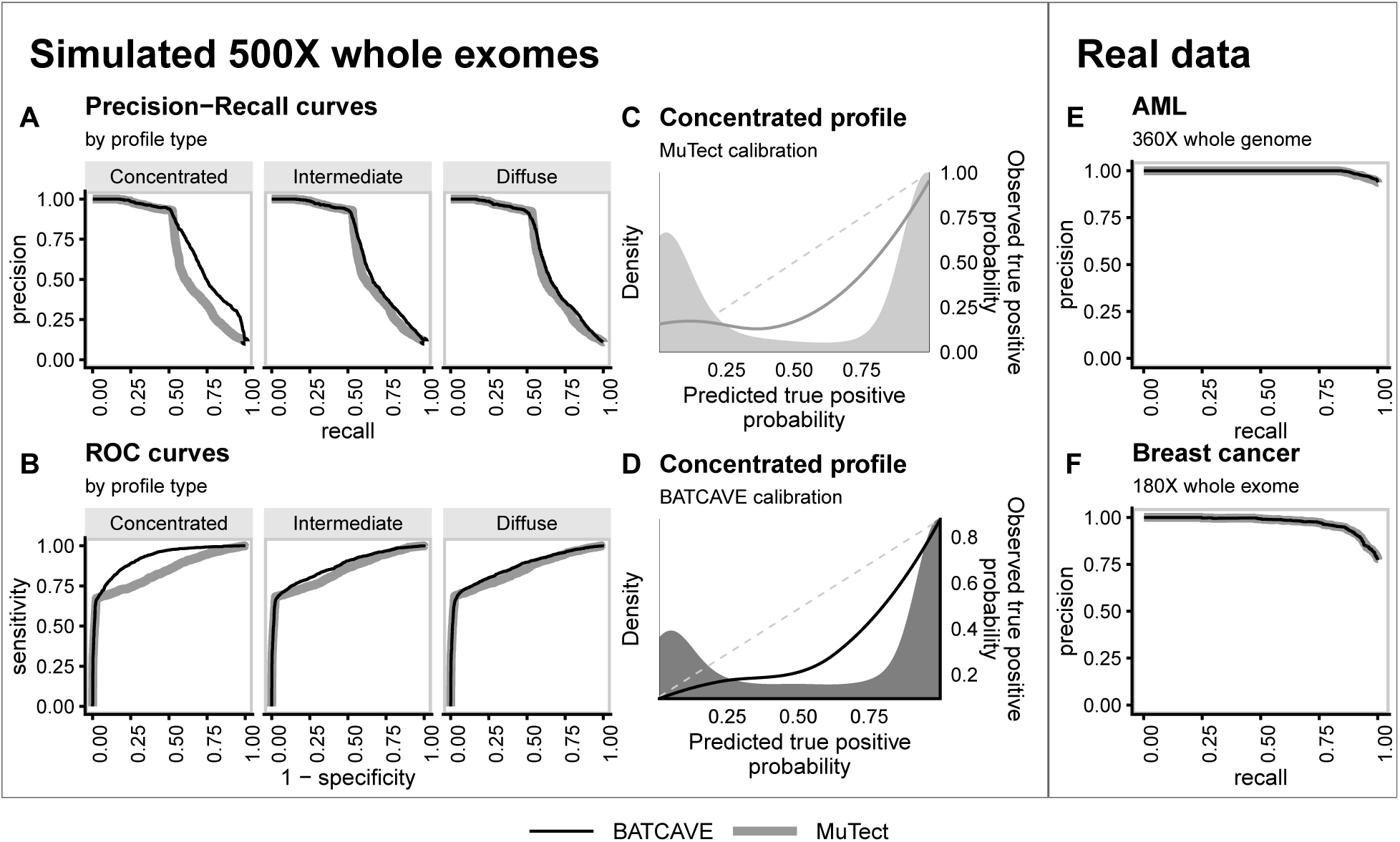
Variant-calling performance on simulated and real data. Throughout, MuTect results are plotted with gray lines and BATCAVE results with black lines. (A) Precision-recall curves and (B) receiver operating characteristic curves for different mutation profiles. (C) and (D) Calibration plots. Shaded regions show distributions of posterior probabilities for true positive variants, and smooth lines show loess-smoothed relationships, from which the Integrated Calibration Index is calculated. For a perfectly calibrated caller, those curves would match the dashed y=x line. (E) and (F) Precision-recall curves for real data in which substantial mutation validation was performed [40, 4].

For all simulated tumors, the estimated mutation rate was approximately 3 · 10^*−*7^ (Table 1), which is lower than the simulated rate of 3 · 10^*−*6^. This is likely due to restrictions within BAMSurgeon, such as sequencing depth and quality, that prevent 100% of simulated variants from being inserted into the reads.

We also assessed calibration, the likelihood that a potential variant with a given posterior probability is actually a true variant. We measured overall calibration performance using the Integrated Calibration Index (ICI) [52], which integrates the difference between predicted and observed probabilities, weighted by the density of the predicted probabilities. This metric is particularly useful in our case, because the density of posterior probabilities is bi-modal (Fig. 4C&D and S1C&D). A large fraction of true negative variants have posterior probabilities less than 10^*−*4^, far below any meaningful threshold, so we evaluated calibration only on potential variants with posterior probability greater than 0.01. For these potential variants, BATCAVE tends to increase posterior probabilities of low probability but true variants (density curves in Fig. 4C&D and S1C&D) while decreasing probabilities of low probability but false variants. For 500X exomes, the calibration of BATCAVE is better than MuTect across the full spectrum of posterior probabilities (Fig. 4 and Table 1). For 100X whole genomes, the calibration of BATCAVE is slightly worse (Fig. S1 and Table 1), likely because there are few low probability true positive variants in tumors sequenced to 100X depth. As with the other metrics, the advantage of BATCAVE increases with the concentration of the mutation profile and the sequencing depth.

In practice, variant callers are typically used with a threshold score above which a variant is called. The user’s choice of threshold ideally meets their need to balance precision and recall; accurate posterior probability estimates enable an informed choice. For posterior probability thresholds between 60 and 90%, the precision of BATCAVE calls is similar to the chosen threshold (Fig. 5&S2). For this range of thresholds, however, the posterior probabilities from MuTect poorly predict precision (Fig. 5&S2). For any posterior probability threshold above 70%, MuTect has a false positive rate of roughly 8%, whereas BATCAVE has a false positive rate that decreases as the threshold increases. The cost of MuTect’s compressed range of posterior probabilities is recall; at any posterior probability threshold BATCAVE has recall better than MuTect. Consequently, BATCAVE posterior probabilities are more informative than MuTect’s with regard to choosing a calling threshold.

**Figure 5:**
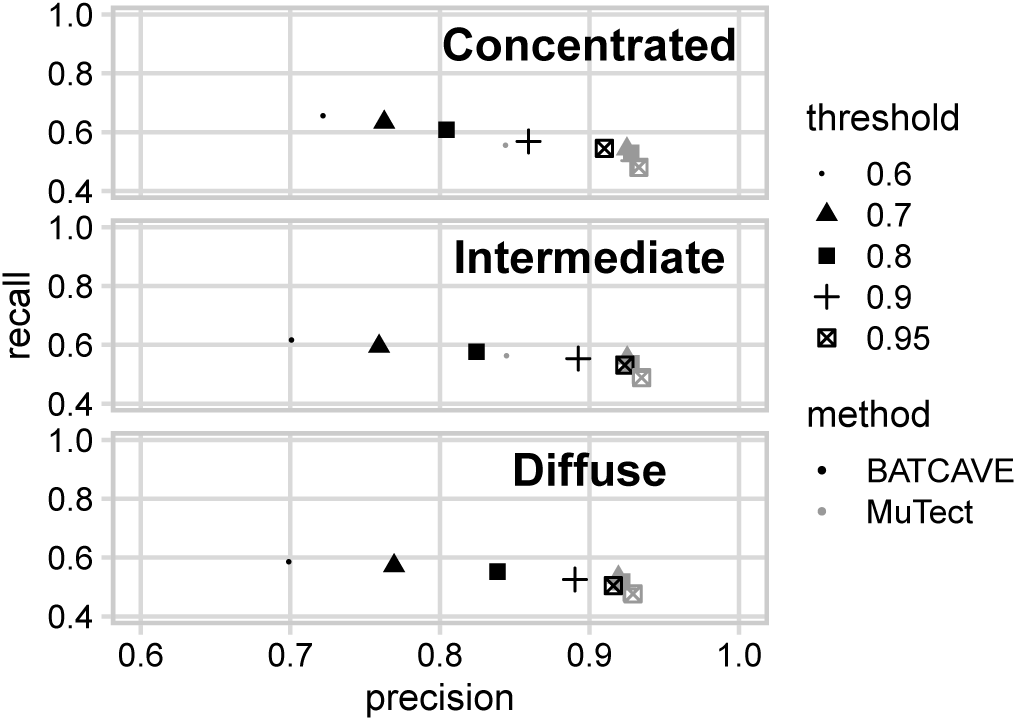
Posterior probability calibration for realistic calling thresholds, for 500X exomes. Plotted is precision and recall for variants identified using various realistic posterior probability thresholds. At these thresholds, the precision of BATCAVE is much closer to the given threshold than MuTect, no matter the concentration of the mutation profile.

### Tests using real tumor data

We tested BATCAVE using two data sets for which deep sequencing and variant validation were performed with the express purpose of evaluating tumor variant calling pipelines, yielding high quality true and false positive data [40, 4]. However, only variants called by at least one variant caller were validated. As a result, there are no validated true or false negative calls, so we considered only precision-recall comparisons for these data.

Griffith et al. sequenced the whole genome of an acute myeloid leukemia (AML) primary tumor to a depth of *>*360X and used targeted sequencing to validate nearly 200,000 mutations [40]. We estimated a per-base mutation rate for this tumor of 4 · 10^*−*8^, which is consistent with previous estimates of AML mutation rates [40, 3]. For both MuTect and BATCAVE, the precision-recall curve is almost perfect for the validated variants (.995 &. 996 area under the curve) (Fig. 4E and Table 1).

Shi et al. performed multi-region whole exome sequencing on six individual breast tumors to a mean target sequencing depth of 160X and validated all variants identified by three different variant calling pipelines [4]. We estimated an average per-base mutation rate for these tumor regions of 4 · 10^*−*8^, which is consistent with observed mutation rates for breast cancers [41] and with the low number of validated somatic mutations. For the validated variants, MuTect and BATCAVE yielded almost identical precision-recall curves (Fig. 4F and Table 1)

## DISCUSSION

BATCAVE is an algorithm that leverages the biology of individual tumor mutation profiles to improve identification of low allelic frequency somatic variants. Our implementation is built on MuTect, one of the most widely used somatic variant callers. BATCAVE improves on the classification accuracy of MuTect in synthetic data (Fig. 4A-D, S1, and Table 1) across the entire range of recall and specificity. Moreover, BATCAVE is better calibrated than MuTect at relevant posterior probability thresholds (Fig. 5 and S2), allowing researchers and clinicians to make informed choices about the trade-off between precision and recall. For real data, testing on validated calls shows that BATCAVE does not degrade performance for variants that are relatively easy to identify (Fig. 4E&F and Table 1). The BATCAVE algorithm can thus be included in a wide variety of sequencing pipelines.

We evaluated BATCAVE with simulated tumors with three different mutation profiles and two real tumors. The simulated diffuse and intermediate profiles (Fig. 2A&B) represent baseline profiles of lung and breast tumors, respectively. And the concentrated profile (Fig. 2C) represents a tumor driven by a particular mutational process, such as UV exposure. But mutational profiles are highly heterogeneous, so concentrated profiles can be found in any tumor type (e.g., Fig. 1C). The two real data sets we considered are among the few for which extensive validation of variant calls has been performed [40, 4]. They happen, however, to have diffuse mutation profiles (Fig. 1A&B), which reduces the expected advantage of BATCAVE over MuTect (Table 1). A more fundamental challenge of using these real data for testing callers is that only a subset of potential variants are validated. This subset tends to be relatively easy to call, so both MuTect and BATCAVE have almost perfect precision and recall for variants that pass heuristic filters (Fig. 4 and Table 1). Moreover, few true negative sites are validated, so specificity and calibration are impossible to calculate. Deep sequencing experiments that validate random samples of uncalled potential variants would give much-needed insight into the differences among statistical models in variant calling.

The improved calibration of BATCAVE posterior probabilities compared to MuTect provides several advantages. In practice, called variants are often manually reviewed to further reduce false positives [53]. Improved calibration enables users to focus review on the most questionable variants. In the clinic, identified variants act as biomarkers for susceptibility to targeted drugs [54]. Well-calibrated posterior probabilities facilitate the use of probabilistic risk models in the choice of treatment [55], rather than an all or nothing approach. For research purposes, the International Cancer Genome Consortium recommends that catalogs of somatic mutations target a precision of 95% and a recall of 80% [56]. Achieving this goal while minimizing cost demands well-calibrated posterior probabilities.

Our current implementation of BATCAVE is as a post-calling algorithm for MuTect, but the algorithm is broadly applicable. We chose to build BATCAVE off MuTect because MuTect is widely used, has state-of-the-art sensitivity and specificity, and includes numerous heuristic filters and alignment adjustments that reduce the prevalence of sequencing errors in results [10, 40]. But the mutational prior can be incorporated into almost any caller with an underlying probabilistic model. For example, Strelka2 computes a joint posterior probability over tumor and normal genotypes, assuming a constant somatic mutation probability at each genomic site [57]. Replacing that constant probability with a mutational prior would require a more complicated manipulation of the quality scores output by Strelka than for MuTect, but it is conceptually straightforward.

The BATCAVE algorithm is computationally inexpensive; our current implementation adds 1 second per 22,000 variants evaluated to a standard GATK best-practices variant calling pipeline. The majority of the computational cost is associated with extracting the trinucleotide context for each potential variant site from the reference genome. Since most callers are already walking the reference genome during the calling process, extracting the trinucleotide context simultaneously would virtually eliminate the computational cost of implementing a mutational prior.

The BATCAVE algorithm incorporates genomic context into the probabilistic model for variant calling. Our current implementation focuses on trinucleotide context, which is known to have a large effect on local mutation rates [58, 59]. There are, however, many other aspects of genomic context that can affect local mutation rates [42], including replication timing [60], expression level [61], and chromatin organization [62]. Some of these, such as replication timing and chromatin organization, could be incorporated into the BATCAVE mutational prior using the empirical distribution of mutations in the human germline [63]. Others, such as expression level, could be tumor-specific, but would require information not available in the variant calls to compute. In the long run, we believe that incorporating more tumor biology into variant calling models will continue to improve performance.

BATCAVE divides the data into two classes: high- and low-confidence variants. The high-confidence variants are used to estimate the mutational prior and mutation rate, which are then used to improve the calling of low-confidence variants. Statistically, this is an empirical Bayesian approach [64], in which the high and low-confidence variants are treated as parallel experiments [65, 66]. In general, high-confidence variants tend to have relatively high allelic frequencies, and consequently tend to have arisen early in tumor development. An implicit assumption of our approach is that the mutational process does not change between high- and low-confidence variants, implying that the mutational profile of the tumor is temporally constant. Recent studies have found differences in mutational profiles among variants of different allelic frequencies [67], although those differences are relatively small. A potential extension of the BATCAVE algorithm is to process potential variants in order of descending allelic frequency and to update the estimated mutational prior as the algorithm proceeds. This approach might increase sensitivity to low-frequency variants generated by recently-arisen mutational processes, at the cost of potentially increasing sensitivity to patterns of sequencing error.

Our results show that adding a mutational prior substantially improves probabilistic variant calling, particularly for tumors with concentrated profiles. Improved variant calling increases the benefit-to-cost ratio of deep sequencing in both research and clinical applications. Moreover, BATCAVE proves to be a better calibrated caller than vanilla MuTect (Fig. 5). Different users will prefer different tradeoffs in terms of precision and recall, which can be more accurately made with BATCAVE. Our R implementation, batcaver, can be easily incorporated into any MuTect-based pipeline, and the mutational profile algorithm can be incorporated into many other callers.

## Software Availability

The `batcaver` R package can be downloaded or installed from http://github.com/bmannakee/batcaver The version of `batcaver` used to generate results and all analysis code have been preserved on Zenodo https://doi.org/10.5281/zenodo.3471715 Python code used to generate simulated tumors has been preserved on Zenodo https://doi.org/10.5281/zenodo.3471741

## ACKNOWLEDGEMENTS

This work was supported by the National Science Foundation via Graduate Research Fellowship award number DGE-1143953 to BKM and by the National Institute of General Medical Sciences of the National Institutes of Health under award number R01GM127348 to RNG. We thank Prof. Edward J. Bedrick for fruitful discussions about the statistical model. This material is based upon High Performance Computing (HPC) resources supported by the University of Arizona TRIF, UITS, and RDI and maintained by the UA Research Technologies department.

---

**Figure S1:**
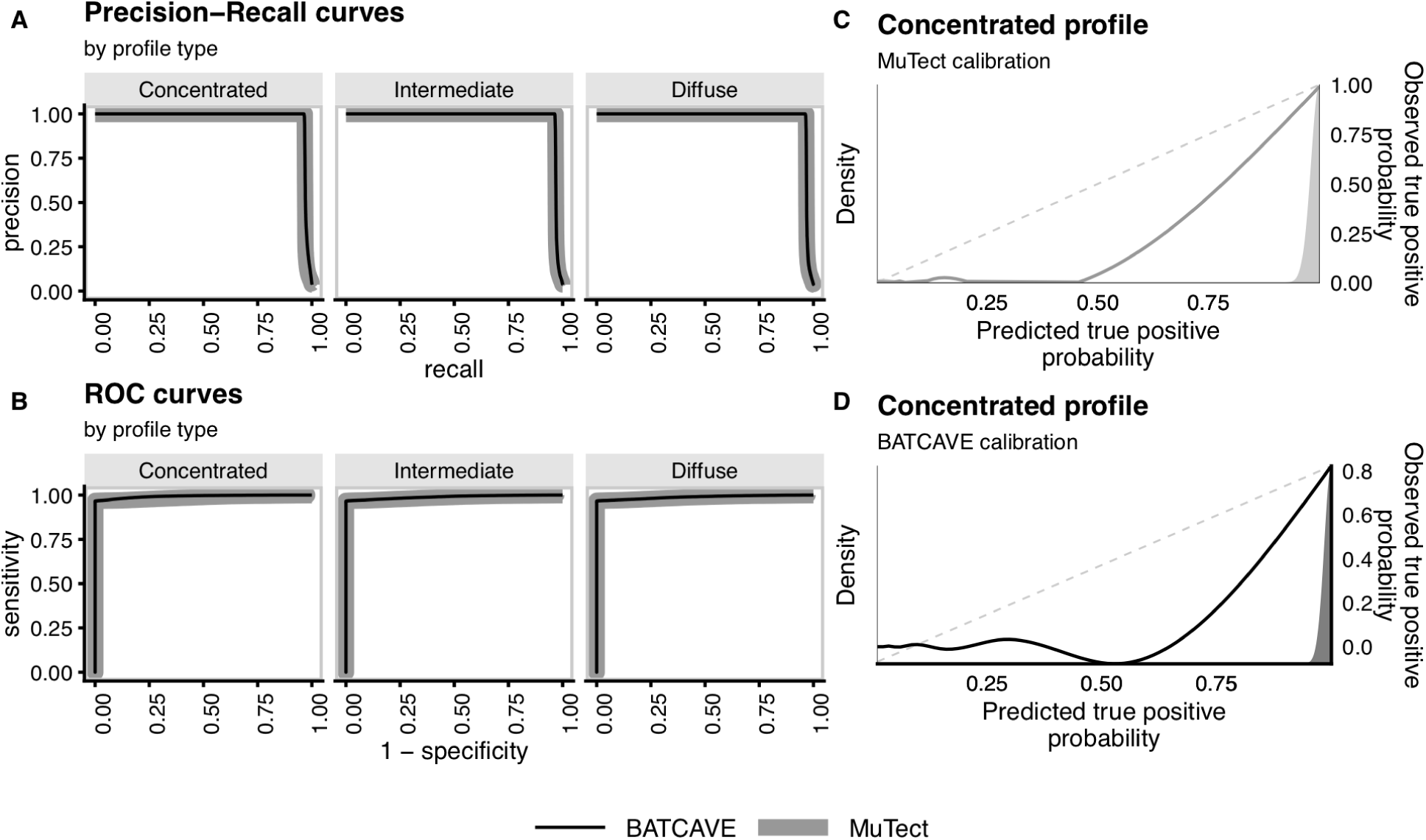
Variant-calling performance on simulated 100X whole genomes. As in Fig. 4A-D, but for 100X whole genomes.

**Figure S2:**
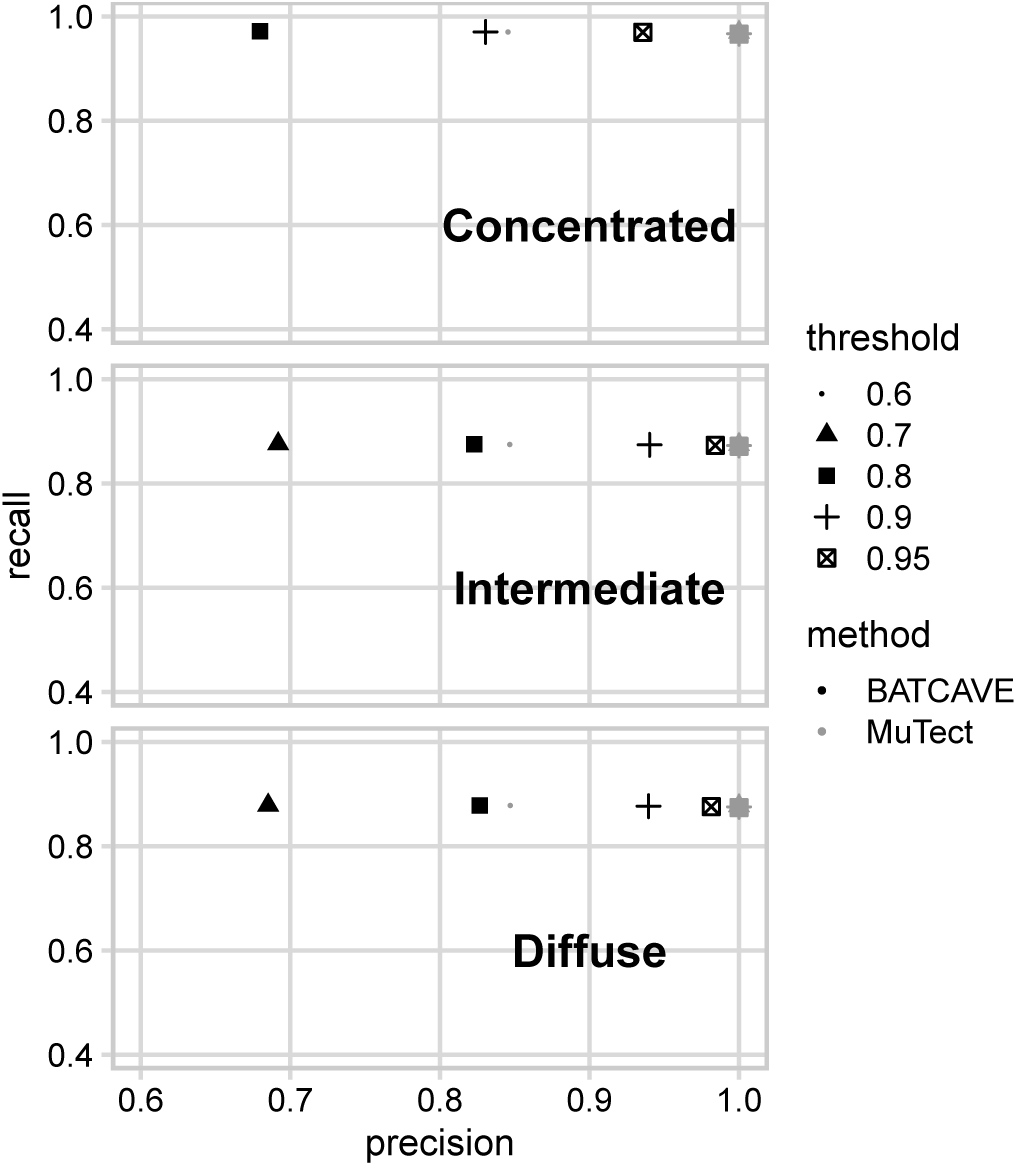
Posterior probability calibration for realistic calling thresholds, for 100X whole genomes. As in Fig. 5, but for 100X whole genomes.

## References

[1] Williams, M. J., Werner, B., Barnes, C. P., Graham, T. A., and Sottoriva, A. (2016) Identification of neutral tumor evolution across cancer types. Nature Genetics, 48(3), 238–244.

[2] Bozic, I., Gerold, J. M., and Nowak, M. A. (2016) Quantifying Clonal and Subclonal Passenger Mutations in Cancer Evolution. PLoS Computational Biology, 12(2), e1004731.

[3] Williams, M. J., Werner, B., Heide, T., Curtis, C., Barnes, C. P., Sottoriva, A., and Graham, T. A. (2018) Quantification of subclonal selection in cancer from bulk sequencing data. Nature Genetics, 50(June), 895–903.

[4] Shi, W., Ng, C. K. Y., Lim, R. S., Pusztai, L., Reis-Filho, J. S., Hatzis, C., Jiang, T., Kumar, S., Li, X., Wali, V. B., Piscuoglio, S., Gerstein, M. B., Chagpar, A. B., and Weigelt, B. (2018) Reliability of Whole-Exome Sequencing for Assessing Intratumor Genetic Heterogeneity I. Cell Reports, 25, 1446–1457.

[5] Ding, J., Bashashati, A., Roth, A., Oloumi, A., Tse, K., Zeng, T., Haffari, G., Hirst, M., Marra, M. A., Condon, A., Aparicio, S., and Shah, S. P. (2012) Feature-based classifiers for somatic mutation detection in tumour-normal paired sequencing data. Bioinformatics, 28(2), 167–175.

[6] Mardis, E. R. (2012) Applying next-generation sequencing to pancreatic cancer treatment. Nature Reviews Gastroenterology & Hepatology, 9(8), 477–486.

[7] Chen, X., Stewart, E., Shelat, A. A., Qu, C., Bahrami, A., Hatley, M., Wu, G., Bradley, C., McEvoy, J., Pappo, A., Spunt, S., Valentine, M. B., Valentine, V., Krafcik, F., Lang, W. H., Wierdl, M., Tsurkan, L., Tolleman, V., Federico, S. M., Morton, C., Lu, C., Ding, L., Easton, J., Rusch, M., Nagahawatte, P., Wang, J., Parker, M., Wei, L., Hedlund, E., Finkelstein, D., Edmonson, M., Shurtleff, S., Boggs, K., Mulder, H., Yergeau, D., Skapek, S., Hawkins, D. S., Ramirez, N., Potter, P. M., Sandoval, J. A., Davidoff, A. M., Mardis, E. R., Wilson, R. K., Zhang, J., Downing, J. R., and Dyer, M. A. (2013) Targeting Oxidative Stress in Embryonal Rhabdomyosarcoma. Cancer Cell, 24(6), 710–724.

[8] Borad, M. J., Champion, M. D., Egan, J. B., Liang, W. S., Fonseca, R., Bryce, A. H., McCullough, A. E., Barrett, M. T., Hunt, K., Patel, M. D., Young, S. W., Collins, J. M., Silva, A. C., Condjella, R. M., Block, M., McWilliams, R. R., Lazaridis, K. N., Klee, E. W., Bible, K. C., Harris, P., Oliver, G. R., Bhavsar, J. D., Nair, A. A., Middha, S., Asmann, Y., Kocher, J. P., Schahl, K., Kipp, B. R., Barr Fritcher, E. G., Baker, A., Aldrich, J., Kurdoglu, A., Izatt, T., Christoforides, A., Cherni, I., Nasser, S., Reiman, R., Phillips, L., McDonald, J., Adkins, J., Mastrian, S. D., Placek, P., Watanabe, A. T., LoBello, J., Han, H., Von Hoff, D., Craig, D. W., Stewart, A. K., and Carpten, J. D. (2014) Integrated Genomic Characterization Reveals Novel, Therapeutically Relevant Drug Targets in FGFR and EGFR Pathways in Sporadic Intrahepatic Cholangiocarcinoma. PLoS Genetics, 10(2).

[9] Findlay, J. M., Castro-Giner, F., Makino, S., Rayner, E., Kartsonaki, C., Cross, W., Kovac, M., Ulahannan, D., Palles, C., Gillies, R. S., Macgregor, T. P., Church, D., Maynard, N. D., Buffa, F., Cazier, J.-B., Graham, T. A., Wang, L.-M., Sharma, R. A., Middleton, M., and Tomlinson, I. (2016) Differential clonal evolution in oesophageal cancers in response to neo-adjuvant chemotherapy. Nature Communications, 7.

[10] Cibulskis, K., Lawrence, M. S., Carter, S. L., Sivachenko, A., Jaffe, D., Sougnez, C., Gabriel, S., Meyerson, M., Lander, E. S., and Getz, G. (2013) Sensitive detection of somatic point mutations in impure and heterogeneous cancer samples. Nature Biotechnology, 31(3), 213–219.

[11] Koboldt, D. C., Zhang, Q., Larson, D. E., Shen, D., McLellan, M. D., Lin, L., Miller, C. A., Mardis, E. R., Ding, L., and Wilson, R. K. (mar, 2012) VarScan 2: Somatic mutation and copy number alteration discovery in cancer by exome sequencing. Genome Research, 22(3), 568–576.

[12] Garrison, E. and Marth, G. (jul, 2012) Haplotype-based variant detection from short-read sequencing. bioRxiv,.

[13] Larson, D. E., Harris, C. C., Chen, K., Koboldt, D. C., Abbott, T. E., Dooling, D. J., Ley, T. J., Mardis, E. R., Wilson, R. K., and Ding, L. (2012) SomaticSniper: identification of somatic point mutations in whole genome sequencing data.. Bioinformatics (Oxford, England), 28(3), 311–7.

[14] Roth, A., Ding, J., Morin, R., Crisan, A., Ha, G., Giuliany, R., Bashashati, A., Hirst, M., Turashvili, G., Oloumi, A., Marra, M. A., Aparicio, S., and Shah, S. P. (2012) JointSNVMix: a probabilistic model for accurate detection of somatic mutations in normal/tumour paired next-generation sequencing data. Bioinformatics, 28(7), 907–913.

[15] Christoforides, A., Carpten, J. D., Weiss, G. J., Demeure, M. J., Von Hoff, D. D., and Craig, D. W. (2013) Identification of somatic mutations in cancer through Bayesian-based analysis of sequenced genome pairs. BMC Genomics, 14, 302.

[16] Jones, D., Raine, K. M., Davies, H., Tarpey, P. S., Butler, A. P., Teague, J. W., Nik-Zainal, S., and Campbell, P. J. (2016) cgpCaVEManWrapper: Simple Execution of CaVEMan in Order to Detect Somatic Single Nucleotide Variants in NGS Data. Current Protocols in Bioinformatics, 56(1), 15.10.1–15.10.18.

[17] Dorri, F., Jewell, S., Bouchard-Côté, A., and Shah, S. P. (2019) Somatic mutation detection and classification through probabilistic integration of clonal population information. Communications Biology, 2(1), 44.

[18] Saunders, C. T., Wong, W. S. W., Swamy, S., Becq, J., Murray, L. J., and Cheetham, R. K. (2012) Strelka: accurate somatic small-variant calling from sequenced tumor-normal sample pairs. Bioinformatics (Oxford, England), 28(14), 1811–7.

[19] Wilm, A., Aw, P. P. K., Bertrand, D., Yeo, G. H. T., Ong, S. H., Wong, C. H., Khor, C. C., Petric, R., Hibberd, M. L., and Nagarajan, N. (2012) LoFreq: A sequence-quality aware, ultra-sensitive variant caller for uncovering cell-population heterogeneity from high-throughput sequencing datasets. Nucleic Acids Research, 40(22), 11189–11201.

[20] Shiraishi, Y., Sato, Y., Chiba, K., Okuno, Y., Nagata, Y., Yoshida, K., Shiba, N., Hayashi, Y., Kume, H., Homma, Y., Sanada, M., Ogawa, S., and Miyano, S. (2013) An empirical Bayesian framework for somatic mutation detection from cancer genome sequencing data. Nucleic Acids Research, 41(7), e89.

[21] Gerstung, M., Beisel, C., Rechsteiner, M., Wild, P., Schraml, P., Moch, H., and Beerenwinkel, N. (2012) Reliable detection of subclonal single-nucleotide variants in tumour cell populations. Nature Communications, 3(May), 811–818.

[22] Carrot-Zhang, J. and Majewski, J. (2017) LoLoPicker: detecting low allelic-fraction variants from low-quality cancer samples. Oncotarget, 8(23), 37032–37040.

[23] Fan, Y., Xi, L., Hughes, D. S., Zhang, J., Zhang, J., Futreal, P. A., Wheeler, D. A., and Wang, W. (2016) MuSE: accounting for tumor heterogeneity using a sample-specific error model improves sensitivity and specificity in mutation calling from sequencing data. Genome biology, 17(1), 178.

[24] Cantarel, B. L., Weaver, D., McNeill, N., Zhang, J., Mackey, A. J., and Reese, J. (apr, 2014) BAYSIC: a Bayesian method for combining sets of genome variants with improved specificity and sensitivity. BMC Bioinformatics, 15(1), 104.

[25] Fang, L. T., Afshar, P. T., Chhibber, A., Mohiyuddin, M., Fan, Y., Mu, J. C., Gibeling, G., Barr, S., Asadi, N. B., Gerstein, M. B., Koboldt, D. C., Wang, W., Wong, W. H., and Lam, H. Y. (2015) An ensemble approach to accurately detect somatic mutations using SomaticSeq. Genome Biology, 16(1), 197.

[26] Spinella, J. F., Mehanna, P., Vidal, R., Saillour, V., Cassart, P., Richer, C., Ouimet, M., Healy, J., and Sinnett, D. (2016) SNooPer: A machine learning-based method for somatic variant identification from low-pass next-generation sequencing. BMC Genomics, 17(1), 912.

[27] Nik-Zainal, S., Alexandrov, L. B., Wedge, D. C., Van Loo, P., Greenman, C. D., Raine, K., Jones, D., Hinton, J., Marshall, J., Stebbings, L. A., Menzies, A., Martin, S., Leung, K., Chen, L., Leroy, C., Ramakrishna, M., Rance, R., Lau, K. W., Mudie, L. J., Varela, I., McBride, D. J., Bignell, G. R., Cooke, S. L., Shlien, A., Gamble, J., Whitmore, I., Maddison, M., Tarpey, P. S., Davies, H. R., Papaemmanuil, E., Stephens, P. J., McLaren, S., Butler, A. P., Teague, J. W., Jönsson, G., Garber, J. E., Silver, D., Miron, P., Fatima, A., Boyault, S., Langerød, A., Tutt, A., Martens, J. W., Aparicio, S. A., Borg, Å., Salomon, A. V., Thomas, G., Børresen-Dale, A.-L., Richardson, A. L., Neuberger, M. S., Futreal, P. A., Campbell, P. J., and Stratton, M. R. (2012) Mutational Processes Molding the Genomes of 21 Breast Cancers. Cell, 149(5), 979–993.

[28] Alexandrov, L. B., Jones, P. H., Wedge, D. C., Sale, J. E., Campbell, P. J., Nik-Zainal, S., and Stratton, M. R. (2015) Clock-like mutational processes in human somatic cells. Nature Genetics, 47(12), 1402–1407.

[29] Lee-Six, H., Øbro, N. F., Shepherd, M. S., Grossmann, S., Dawson, K., Belmonte, M., Osborne, R. J., Huntly, B. J. P., Martincorena, I., Anderson, E., O’Neill, L., Stratton, M. R., Laurenti, E., Green, A. R., Kent, D. G., and Campbell, P. J. (sep, 2018) Population dynamics of normal human blood inferred from somatic mutations. Nature, 561(7724), 473–478.

[30] Alexandrov, L. B., Nik-Zainal, S., Wedge, D. C., Campbell, P. J., and Stratton, M. R. (2013) Deciphering Signatures of Mutational Processes Operative in Human Cancer. Cell Reports, 3(1), 246–259.

[31] Helleday, T., Eshtad, S., and Nik-Zainal, S. (jul, 2014) Mechanisms underlying mutational signatures in human cancers. Nature Reviews Genetics, 15(9), 585–598.

[32] Nik-Zainal, S., Davies, H., Staaf, J., Ramakrishna, M., Glodzik, D., Zou, X., Martincorena, I., Alexandrov, L. B., Martin, S., Wedge, D. C., Van Loo, P., Ju, Y. S., Smid, M., Brinkman, A. B., Morganella, S., Aure, M. R., Lingjærde, O. C., Langerød, A., Ringnér, M., Ahn, S.-M., Boyault, S., Brock, J. E., Broeks, A., Butler, A., Desmedt, C., Dirix, L., Dronov, S., Fatima, A., Foekens, J. A., Gerstung, M., Hooijer, G. K. J., Jang, S. J., Jones, D. R., Kim, H.-Y., King, T. A., Krishnamurthy, S., Lee, H. J., Lee, J.-Y., Li, Y., McLaren, S., Menzies, A., Mustonen, V., O’Meara, S., Pauporté, I., Pivot, X., Purdie, C. A., Raine, K., Ramakrishnan, K., Rodríguez-González, F. G., Romieu, G., Sieuwerts, A. M., Simpson, P. T., Shepherd, R., Stebbings, L., Stefansson, O. A., Teague, J., Tommasi, S., Treilleux, I., Van den Eynden, G. G., Vermeulen, P., Vincent-Salomon, A., Yates, L., Caldas, C., van’t Veer, L., Tutt, A., Knappskog, S., Tan, B. K. T., Jonkers, J., Borg, Å., Ueno, N. T., Sotiriou, C., Viari, A., Futreal, P. A., Campbell, P. J., Span, P. N., Van Laere, S., Lakhani, S. R., Eyfjord, J. E., Thompson, A. M., Birney, E., Stunnenberg, H. G., van de Vijver, M. J., Martens, J. W. M., Børresen-Dale, A.-L., Richardson, A. L., Kong, G., Thomas, G., and Stratton, M. R. (2016) Landscape of somatic mutations in 560 breast cancer whole-genome sequences. Nature, 534(7605), 47–54.

[33] Kandoth, C., McLellan, M. D., Vandin, F., Ye, K., Niu, B., Lu, C., Xie, M., Zhang, Q., McMichael, J. F., Wyczalkowski, M. A., Leiserson, M. D. M., Miller, C. A., Welch, J. S., Walter, M. J., Wendl, M. C., Ley, T. J., Wilson, R. K., Raphael, B. J., and Ding, L. (2013) Mutational landscape and significance across 12 major cancer types. Nature, 502(7471), 333–339.

[34] Alexandrov, L. B., Ju, Y. S., Haase, K., Van Loo, P., Martincorena, I., Nik-Zainal, S., Totoki, Y., Fujimoto, A., Nakagawa, H., Shibata, T., Campbell, P. J., Vineis, P., Phillips, D. H., and Stratton, M. R. (2016) Mutational signatures associated with tobacco smoking in human cancer. Science, 354(6312), 618–622.

[35] Stephens, P., Edkins, S., Davies, H., Greenman, C., Cox, C., Hunter, C., Bignell, G., Teague, J., Smith, R., Stevens, C., O’Meara, S., Parker, A., Tarpey, P., Avis, T., Barthorpe, A., Brackenbury, L., Buck, G., Butler, A., Clements, J., Cole, J., Dicks, E., Edwards, K., Forbes, S., Gorton, M., Gray, K., Halliday, K., Harrison, R., Hills, K., Hinton, J., Jones, D., Kosmidou, V., Laman, R., Lugg, R., Menzies, A., Perry, J., Petty, R., Raine, K., Shepherd, R., Small, A., Solomon, H., Stephens, Y., Tofts, C., Varian, J., Webb, A., West, S., Widaa, S., Yates, A., Brasseur, F., Cooper, C. S., Flanagan, A. M., Green, A., Knowles, M., Leung, S. Y., Looijenga, L. H. J., Malkowicz, B., Pierotti, M. A., Teh, B., Yuen, S. T., Nicholson, A. G., Lakhani, S., Easton, D. F., Weber, B. L., Stratton, M. R., Futreal, P. A., and Wooster, R. (2005) A screen of the complete protein kinase gene family identifies diverse patterns of somatic mutations in human breast cancer. Nature Genetics, 37(6), 590–592.

[36] Burrell, R. A., McGranahan, N., Bartek, J., and Swanton, C. (2013) The causes and consequences of genetic heterogeneity in cancer evolution. Nature, 501(7467), 338–345.

[37] Nakamura, H., Arai, Y., Totoki, Y., Shirota, T., Elzawahry, A., Kato, M., Hama, N., Hosoda, F., Urushidate, T., Ohashi, S., Hiraoka, N., Ojima, H., Shimada, K., Okusaka, T., Kosuge, T., Miyagawa, S., and Shibata, T. (2015) Genomic spectra of biliary tract cancer. Nature Genetics, 47(9), 1003–1010.

[38] Witkiewicz, A. K., McMillan, E. A., Balaji, U., Baek, G., Lin, W.-C., Mansour, J., Mollaee, M., Wagner, K.-U., Koduru, P., Yopp, A., Choti, M. A., Yeo, C. J., McCue, P., White, M. A., and Knudsen, E. S. (2015) Whole-exome sequencing of pancreatic cancer defines genetic diversity and therapeutic targets. Nature Communications, 6, 6744.

[39] Kumar, A., Coleman, I., Morrissey, C., Zhang, X., True, L. D., Gulati, R., Etzioni, R., Bolouri, H., Montgomery, B., White, T., Lucas, J. M., Brown, L. G., Dumpit, R. F., DeSarkar, N., Higano, C., Yu, E. Y., Coleman, R., Schultz, N., Fang, M., Lange, P. H., Shendure, J., Vessella, R. L., and Nelson, P. S. (2016) Substantial interindividual and limited intraindividual genomic diversity among tumors from men with metastatic prostate cancer. Nature Medicine, 22(4), 369–378.

[40] Griffith, M., Miller, C. A., Griffith, O. L., Krysiak, K., Skidmore, Z. L., Ramu, A., Walker, J. R., Dang, H. X., Trani, L., Larson, D. E., Demeter, R. T., Wendl, M. C., McMichael, J. F., Austin, R. E., Magrini, V., McGrath, S. D., Ly, A., Kulkarni, S., Cordes, M. G., Fronick, C. C., Fulton, R. S., Maher, C. A., Ding, L., Klco, J. M., Mardis, E. R., Ley, T. J., and Wilson, R. K. (2015) Optimizing Cancer Genome Sequencing and Analysis. Cell Systems, 1(3), 210–223.

[41] Alexandrov, L. B., Kim, J., Haradhvala, N. J., Huang, M. N., Ng, A. W., Wu, Y., Boot, A., Covington, K. R., Gordenin, D. A., Bergstrom, E. N., Islam, S. M. A., Lopez-Bigas, N., Klimczak, L. J., McPherson, J. R., Morganella, S., Sabarinathan, R., Wheeler, D. A., Mustonen, V., Group, t. P. M. S. W., Getz, G., Rozen, S. G., and Stratton, M. R. (2019) The Repertoire of Mutational Signatures in Human Cancer. bioRxiv, p. 322859.

[42] Buisson, R., Langenbucher, A., Bowen, D., Kwan, E. E., Benes, C. H., Zou, L., and Lawrence, M. S. (2019) Passenger hotspot mutations in cancer driven by APOBEC3A and mesoscale genomic features.. Science (New York, N.Y.), 364(6447), eaaw2872.

[43] Pagès, H. BSgenome: Software infrastructure for efficient representation of full genomes and their SNPs. (2019).

[44] Lawrence, M., Huber, W., Pagès, H., Aboyoun, P., Carlson, M., Gentleman, R., Morgan, M., and Carey, V. (2013) Software for Computing and Annotating Genomic Ranges. PLoS Computational Biology, 9(8).

[45] Obenchain, V., Lawrence, M., Carey, V., Gogarten, S., Shannon, P., and Morgan, M. (2014) VariantAnnotation: a Bioconductor package for exploration and annotation of genetic variants. Bioinformatics, 30(14), 2076–2078.

[46] Gehring, J. S., Fischer, B., Lawrence, M., and Huber, W. (2015) SomaticSignatures: inferring mutational signatures from single-nucleotide variants. Bioinformatics, 31(22), 3673–3675.

[47] COSMIC Mutational Signatures (Version 2) https://cancer.sanger.ac.uk/cosmic/signatures_v2.

[48] Alexandrov, L. B., Nik-Zainal, S., Wedge, D. C., Aparicio, S. A. J. R., Behjati, S., Biankin, A. V., Bignell, G. R., Bolli, N., Borg, A., Børresen-Dale, A.-L., Boyault, S., Burkhardt, B., Butler, A. P., Caldas, C., Davies, H. R., Desmedt, C., Eils, R., Eyfjörd, J. E., Foekens, J. A., Greaves, M., Hosoda, F., Hutter, B., Ilicic, T., Imbeaud, S., Imielinsk, M., Jäger, N., Jones, D. T. W., Jones, D., Knappskog, S., Kool, M., Lakhani, S. R., López-Otín, C., Martin, S., Munshi, N. C., Nakamura, H., Northcott, P. A., Pajic, M., Papaemmanuil, E., Paradiso, A., Pearson, J. V., Puente, X. S., Raine, K., Ramakrishna, M., Richardson, A. L., Richter, J., Rosenstiel, P., Schlesner, M., Schumacher, T. N., Span, P. N., Teague, J. W., Totoki, Y., Tutt, A. N. J., Valdés-Mas, R., van Buuren, M. M., van ’t Veer, L., Vincent-Salomon, A., Waddell, N., Yates, L. R., Zucman-Rossi, J., Andrew Futreal, P., McDermott, U., Lichter, P., Meyerson, M., Grimmond, S. M., Siebert, R., Campo, E., Shibata, T., Pfister, S. M., Campbell, P. J., and Stratton, M. R. (2013) Signatures of mutational processes in human cancer. Nature, 500(7463), 415–421.

[49] Mu, J. C., Mohiyuddin, M., Li, J., Bani Asadi, N., Gerstein, M. B., Abyzov, A., Wong, W. H., and Lam, H. Y. K. (2015) VarSim: a high-fidelity simulation and validation framework for high-throughput genome sequencing with cancer applications. Bioinformatics, 31(9), 1469–1471.

[50] Li, H. and Durbin, R. (2009) Fast and accurate short read alignment with Burrows-Wheeler transform. Bioinformatics, 25(14), 1754–1760.

[51] Ewing, A. D., Houlahan, K. E., Hu, Y., Ellrott, K., Caloian, C., Yamaguchi, T. N., Bare, J. C., P’ng, C., Waggott, D., Sabelnykova, V. Y., Kellen, M. R., Norman, T. C., Haussler, D., Friend, S. H., Stolovitzky, G., Margolin, A. A., Stuart, J. M., Boutros, P. C., Li, C., Bertrand, D., Nagarajan, N., Chen, Q.-R., Hsu, C.-H., Hu, Y., Yan, C., Kibbe, W., Meerzaman, D., Cibulskis, K., Rosenberg, M., Bergelson, L., Kiezun, A., Radenbaugh, A., Sertier, A.-S., Ferrari, A., Tonton, L., Bhutani, K., Hansen, N. F., Wang, D., Song, L., Lai, Z., Liao, Y., Shi, W., Carbonell-Caballero, J., Dopazo, J., Lau, C. C. K., Guinney, J., Kellen, M. R., Norman, T. C., Haussler, D., Friend, S. H., Stolovitzky, G., Margolin, A. A., Stuart, J. M., and Boutros, P. C. (2015) Combining tumor genome simulation with crowdsourcing to benchmark somatic single-nucleotide-variant detection. Nature Methods, 12(7), 623–630.

[52] Austin, P. C. and Steyerberg, E. W. (2019) The Integrated Calibration Index (ICI) and related metrics for quantifying the calibration of logistic regression models. Statistics in Medicine, p. sim.8281.

[53] Barnell, E. K., Ronning, P., Campbell, K. M., Krysiak, K., Ainscough, B. J., Sheta, L. M., Pema, S. P., Schmidt, A. D., Richters, M., Cotto, K. C., Danos, A. M., Ramirez, C., Skidmore, Z. L., Spies, N. C., Hundal, J., Sediqzad, M. S., Kunisaki, J., Gomez, F., Trani, L., Matlock, M., Wagner, A. H., Swamidass, S. J., Griffith, M., and Griffith, O. L. (2019) Standard operating procedure for somatic variant refinement of sequencing data with paired tumor and normal samples. Genetics in Medicine, 21(4), 972–981.

[54] Boutros, P. C. (2015) The path to routine use of genomic biomarkers in the cancer clinic.. Genome research, 25(10), 1508–13.

[55] Holmberg, L. and Vickers, A. (2013) Evaluation of Prediction Models for Decision-Making: Beyond Calibration and Discrimination. PLoS Medicine, 10(7), e1001491.

[56] International Cancer Genome Consortium Goals, Structure, Policies, and Guidelines https://icgc.org/icgc/goals-structure-policies-guidelines/e8-genome-analyses.

[57] Kim, S., Scheffler, K., Halpern, A. L., Bekritsky, M. A., Noh, E., Källberg, M., Chen, X., Kim, Y., Beyter, D., Krusche, P., and Saunders, C. T. (2018) Strelka2: fast and accurate calling of germline and somatic variants. Nature Methods, 15(8), 591–594.

[58] Martincorena, I. and Campbell, P. J. (2015) Somatic mutation in cancer and normal cells.. Science (New York, N.Y.), 349(6255), 1483–9.

[59] Hollstein, M., Alexandrov, L. B., Wild, C. P., Ardin, M., and Zavadil, J. (2017) Base changes in tumour DNA have the power to reveal the causes and evolution of cancer. Oncogene, 36(2), 158–167.

[60] Stamatoyannopoulos, J. A., Adzhubei, I., Thurman, R. E., Kryukov, G. V., Mirkin, S. M., and Sunyaev, S. R. (apr, 2009) Human mutation rate associated with DNA replication timing. Nature Genetics, 41(4), 393–395.

[61] Pleasance, E. D., Cheetham, R. K., Stephens, P. J., McBride, D. J., Humphray, S. J., Greenman, C. D., Varela, I., Lin, M.-L., Ordóñez, G. R., Bignell, G. R., Ye, K., Alipaz, J., Bauer, M. J., Beare, D., Butler, A., Carter, R. J., Chen, L., Cox, A. J., Edkins, S., Kokko-Gonzales, P. I., Gormley, N. A., Grocock, R. J., Haudenschild, C. D., Hims, M. M., James, T., Jia, M., Kingsbury, Z., Leroy, C., Marshall, J., Menzies, A., Mudie, L. J., Ning, Z., Royce, T., Schulz-Trieglaff, O. B., Spiridou, A., Stebbings, L. A., Szajkowski, L., Teague, J., Williamson, D., Chin, L., Ross, M. T., Campbell, P. J., Bentley, D. R., Futreal, P. A., and Stratton, M. R. (2010) A comprehensive catalogue of somatic mutations from a human cancer genome. Nature, 463(7278), 191–196.

[62] Schuster-Böckler, B. and Lehner, B. (2012) Chromatin organization is a major influence on regional mutation rates in human cancer cells. Nature, 488(7412), 504–507.

[63] Hodgkinson, A. and Eyre-Walker, A. (2011) Variation in the mutation rate across mammalian genomes. Nature Reviews Genetics, 12(11), 756–766.

[64] Robbins, H. (1954) An empirical Bayes approach to statistics. In Proceedings of the Third Berkeley Symposium on Mathematical Statistics and Probability Vol. 1, pp. 157–163.

[65] Morris, C. N. (1983) Parametric Empirical Bayes Inference: Theory and Applications. Journal of the American Statistical Association, 78, 47–55.

[66] Efron, B. (2014) Two modeling strategies for empirical Bayes estimation. Statistical science : a review journal of the Institute of Mathematical Statistics, 29(2), 285–301.

[67] Rubanova, Y., Shi, R., Li, R., Wintersinger, J., Deshwar, A., Sahin, N., and Morris, Q. (2018) Reconstructing Evolutionary Trajectories of Mutations in Cancer. bioRxiv,.

